# The clock-modulatory activity of Nobiletin suppresses adipogenesis via Wnt signaling

**DOI:** 10.1101/2023.02.07.527587

**Authors:** Xuekai Xiong, Tali Kiperman, Weini Li, Sangeeta Dhawan, Jeongkyung Lee, Vijay Yechoor, Ke Ma

**Author notes:** These authors contributed equally to the manuscript (Tali Kiperman, Weini Li).

## Abstract

The circadian clock machinery exerts transcriptional control to modulate adipogenesis and its disruption leads to the development of obesity. Here we report that Nobiletin, a clock amplitude-enhancing molecule, displays anti-adipogenic properties via activating a clock-controlled Wnt signaling pathway that suppresses adipocyte differentiation. Nobiletin augmented clock oscillation with period length shortening in the adipogenic mesenchymal precursor cells and preadipocytes, accompanied by an induction of Bmal1 and core clock components. Consistent with its circadian clock-modulatory activity, Nobiletin inhibited the lineage commitment and terminal differentiation of adipogenic progenitors. Mechanistically, we show that Nobiletin induced the re-activation of Wnt signaling during adipogenic differentiation via transcriptional up-regulation of key components of this pathway. Furthermore, Nobiletin administration in mice markedly reduced adipocyte hypertrophy, leading to a significant loss of fat mass and body weight reduction. Lastly, Nobiletin inhibited the maturation of primary preadipocytes and this effect was dependent on a functional clock regulation. Collectively, our findings uncover a novel activity of Nobiletin in suppressing adipocyte development, implicating its potential therapeutic application in countering obesity and its associated metabolic consequences.

## Introduction

The circadian clock, a time-keeping mechanism that drives rhythmic oscillations of biological processes, exerts pervasive control in metabolic regulation (Bass and Takahashi, 2010; Takahashi, 2017). The rate-limiting enzymes involved in glucose metabolism, mitochondrial oxidative phosphorylation, and lipid utilization display a circadian clock transcriptional control (Panda et al., 2002), along with diurnal orchestration of global metabolome evident for metabolic substrates (Eckel-Mahan et al., 2012). Disruption of this timing mechanism, increasingly prevalent in a modern lifestyle, predisposes to metabolic dysregulations leading to the development of obesity and diabetes (Bass and Takahashi, 2010; Stenvers et al., 2019). Maintaining or re-enforcing circadian clock control to augment the temporal coordination of its metabolic outputs may provide benefits for the prevention or treatment of metabolic diseases (Cederroth et al., 2019; Hatori et al., 2012; Sulli et al., 2018). There is increasing interests in identifying clock-modulating molecules that may yield metabolic benefits for combating obesity, type II diabetes and related complications (Chen et al., 2018).

The circadian clock is driven by a negative molecular feedback loop consisting of transcriptional and translational regulations (Takahashi, 2017). Key molecular clock transcription factors, CLOCK (Circadian Locomotor Output Cycles Kaput) and its heterodimer partner Bmal1, activate clock gene transcription. Direct targets of CLOCK/Bmal1, transcriptional repressors including the Period and Cryptochrome genes, constitute the negative feed-back limb that represses CLOCK/Bmal1 activity. The adipose tissue possesses tissue-intrinsic peripheral clocks, with cell-autonomous clock oscillations observed in adipogenic progenitors and mature adipocytes (Otway et al., 2009; Zvonic et al., 2006). In preadipocytes, daily rhythms of commitment to differentiation occur, driven by an accumulation of adipogenic factors (Zhang et al., 2022). Accumulating evidence indicates that circadian clock dysregulation is intimately linked with the development of obesity (Karatsoreos et al., 2011; Kolbe et al., 2019). Clock disruption via a shiftwork regimen leads to adipose tissue expansion with inflammation and tissue fibrosis, significant contributors to systemic insulin resistance (Xiong et al., 2021). Conversely, time-restricted feeding that augments clock regulation prevents high-fat diet induced obesity (Hatori et al., 2012), and Rev-erbα activation by synthetic agonists resulted in anti-obesity and anti-lipidemic effects (Solt et al., 2012). Thus, pharmacological interventions targeting clock modulation to maintain or augment circadian control in adipose tissue may have anti-obesity therapeutic potentials.

Various components of the molecular clock machinery, a transcriptional-translation feedback loop, are involved in adipocyte biology (Nam et al., 2016). The essential clock activator Bmal1 exerts direct translational control of the Wnt pathway to inhibit adipogenesis, with its loss of function resulting in obesity (Guo et al., 2012). Similarly, circadian control of the actin cytoskeleton-SRF/MRTF regulatory cascade suppresses beige adipocyte development (Xiong et al., 2022a). In contrast, Rev-erbα, a direct target of Bmal1 and a transcriptional repressor of the clock regulatory loop, is required for brown adipogenesis (Nam et al., 2015a). In addition, Period proteins, including Per2 and Per3 that constitute the repressive feedback arm of the clock, are involved in adipocyte precursor development into adipocytes (Aggarwal et al., 2017; Grimaldi et al., 2010). Retinoid-related orphan receptors, RORα/γ, are positive clock regulators that drive gene activation via a RORE DNA-binding regulatory element (Jetten, 2009). RORs can be induced by adipogenic differentiation (Austin et al., 1998), and RORα was found to inhibit mature adipocyte differentiation by interfering with the activity of adipogenic factor C/EBPß that blocks PPARγ and C/EBPα induction of terminal differentiation (Ohoka et al., 2009). Interestingly, RORα deficiency in the staggerer mice (Rorα^sg^) due to a spontaneous deletion mutation promotes thermogenic induction of brown and beige adipocytes (Lau et al., 2015).

Nobiletin (Nob), a flavonoid compound found in citrus fruits, is known to display beneficial metabolic properties (Assini et al., 2013). A recent study revealed the agonist activity of Nob for RORα/γ, key clock components (He et al., 2016). Nob thus functions as a clock modulator that augments circadian oscillatory amplitude due to RORα/γ activation with consequent induction of Bmal1 transcription. Despite previous reports of Nob effect on promoting skeletal muscle mitochondrial oxidation and its lipid-lowering action (Assini et al., 2013; Nohara et al., 2019), whether it directly modulates cell-autonomous adipocyte clock to impact adipocyte function remains unknown. Based on the role of circadian clock in orchestrating adipogenic differentiation (Guo et al., 2012; Nam et al., 2015a; Xiong et al., 2022a; Zhang et al., 2022), we tested whether Nob could influence the development of adipogenic progenitors to mature adipocytes. Using distinct adipogenic cellular models together with *in vivo* testing, the current study uncovered a novel clock-dependent action of Nob in augmenting Wnt signaling to suppress adipocyte formation.

## Materials & Methods

### Animals

Mice were maintained in the City of Hope vivarium under a constant 12:12 light dark cycle, with lights on at 6:00 AM (ZT0). All animal experiments were approved by the Institutional Animal Care & Use Committee (IACUC) of City of Hope and carried out in concordance with the IACUC approval. C57BL/6J mice were purchased from Jackson Laboratory and used for experiments following 2 weeks or longer of acclimation at 10 or 40 weeks of age. Nobiletin for 0.1% and 0.2% pellet diets was obtained from Hangzhou R&S Pharmchem Co and used for custom diet synthesis by Research Diets together with chow control diet.

### Cell culture and adipogenic differentiation

3T3-L1 and C3H10T1/2 were obtained from ATCC, and maintained in DMEM with 10% fetal bovine serum supplemented with Penicillin-Streptomycin-Glutamine, as previously described (Liu et al., 2020; Nam et al., 2015b). For adipogenic differentiation of these cell lines, induction media (1.6μM insulin, 1μM dexamethasone, 0.5mM IBMX, 0.5 uM Rosiglitazone) was used for 3 days followed by maintenance medium with insulin for 3 days for 3T3-L1 and 5 days for C3H10T1/2 cells, as previously described (Liu et al., 2020). Nobiletin used for cell culture was purchased from Cayman Chemicals.

### Generation of stable adipogenic progenitor cell lines containing Per2::dLuc luciferase reporter

3T3-L1 preadipocytes and C3H10T1/2 mesenchymal stem cells obtained from ATCC were used for lentiviral transduction containing a *Per2::dLuc* luciferase reporter (Yoo et al., 2004), with stable clone selection using puromycin, as described previously (Guo et al., 2012). Briefly, cells were transfected with lentiviral packaging plasmids (pSPAX.2 and pMD2.G) and lentivirus vector *Per2::dLuc* using PEI Max (Polysciences). At 48 hours post-transfection, lentiviruses were collected. 3T3-L1 and C3H10T1/2 cells were infected using collected lentiviral media supplemented with polybrene. 24 hours following lentiviral infection, stable cell lines were selected in the presence of 2 μg/ml puromycin.

### Primary preadipocyte isolation and adipogenic induction

The stromal vascular fraction containing preadipocytes were isolated from subcutaneous fat pads, as described (Xiong et al., 2022a). Briefly, fat pads were cut into pieces and digested using 0.1% collagenase Type 1 with 0.8% BSA at 37^0^C in a horizontal shaker for 60 minutes, passed through Nylon mesh and centrifuged to collect the pellet containing the stromal vascular fraction with preadipocytes. Preadipocytes were cultured in F12/DMEM supplemented with bFGF (2.5 ng/ml), expanded for two passages and subjected to differentiation in 6-well plates at 90% confluency. Adipogenic differentiation was induced for 2 days in medium containing 10% FBS, 1.6 μM insulin, 1 μM dexamethasone, 0.5 mM IBMX, 0.5 uM rosiglitazone before switching to maintenance medium for 4 days with insulin only. Nobiletin at indicated concentrations were added for the entire differentiation time course.

### Oil-red-O and Bodipy staining

These staining for neutral lipids during adipogenic differentiation were performed as previously described (Nam et al., 2015b). Briefly, for oil-red-O staining, cells were fixed using 10% formalin and incubated using 0.5% oil-red-O solution for 1 hour. Bodipy 493/503 was used at 1mg/L together with DAPI for 15 minutes, following 4% paraformaldehyde fixation and permeabilization with 0.2% triton-X100.

### Continuous Bioluminescence monitoring of Per2::dLuc luciferase reporter

3T3-L1 and C3H10T1/2 cells containing a *Per2::dLuc* luciferase reporter were seeded at 4×10^5^ density on 24 well plates and real-time luciferase activity recorded for 7 days using LumiCycle 92 (ActiMetrics), as previously described (Xiong et al., 2022b). Briefly, cells were used at 90% confluence following overnight culture with explant medium luciferase recording media. Explant medium contains 50% 2xDMEM buffer stock, 10% FBS (Cytiva), 1% PSG, pH7 1M HEPES, 7.5% Sodium Bicarbonate, sodium hydroxide (100 mM) and XenoLight D-Luciferin bioluminiscent substrate (100 mM). Raw and subtracted results of real-time bioluminescence recording data for 6 days were exported, and data was calculated as luminescence counts per second. LumiCycle Analysis Program (Actimetrics) was used to determine clock oscillation period, length amplitude and phase. Briefly, raw data following the first cycle from day 2 to day 5 were fitted to a linear baseline, and the baseline-subtracted data (polynomial number = 1) were fitted to a sine wave, from which period length and goodness of fit and damping constant were determined. For samples that showed persistent rhythms, goodness-of-fit of >80% was usually achieved.

### TOPFlash luciferase reporter assay

M50 Super 8xTOPFlash luciferase reporter containing Wnt-responsive TCF bindings sites was a gift from Randall Moon (Veeman et al., 2003) obtained from Addgene (Addgene plasmid # 12456) TOPFlash. Cells were seeded and grown overnight. At 90% confluence cell transfection was performed. 24 hours following transfection, cells were treated using 10% Wnt3a conditioned media obtained from L Wnt-3A cell line (ATCC CRL-2647) to induce luciferase activity. Luciferase activity was assayed using Dual-Luciferase Reporter Assay Kit (Promega) in 96-well black plates. TOPFlash luciferase reporter luminescence was measured on microplate reader (TECAN infinite M200pro) and normalized to control FOPFlash activity, as previously described (Guo et al., 2012). The mean and standard deviation values were calculated for each well and graphed.

### Immunoblot analysis

Protein was extracted using lysis buffer (3% NaCl, 5% Tris-HCl, 10% Glycerol, 0.5% Triton X-10 in Sterile MilliQ water) containing protease inhibitor. 20-40 μg of total protein was resolved on 10% SDS-PAGE gels followed by immunoblotting on PVDF membranes (Bio-rad). Membranes were developed by chemiluminescence (SuperSignal West Pico, Pierce Biotechnology) and signals were obtained via a chemiluminescence imager (Amersham Imager 680, GE Biosciences). Antibodies used are listed in Supplemental Table 1.

### RNA extraction and RT-qPCR analysis

Trizol (Invitrogen) and PureLink RNA Mini Kit (Invitrogen) were used to isolate total RNA from snap-frozen tissues or cells, respectively. cDNA was generated using Revert Aid RT kit (ThermoFisher) and quantitative PCR was performed using SYBR Green Master Mix (Thermo Fisher) in triplicates on ViiA 7 Real-Time PCR System (Applied Biosystems). Relative expression levels were determined using the comparative Ct method with normalization to 36B4 as internal control. PCR primers sequence are listed in Supplemental Table 2.

### Hematoxylin/eosin histology and adipocyte size distribution

Adipose tissues were fixed with 10% neutral-buffered formalin for 72 hours prior to embedding. 10μm paraffin sections were processed for hematoxylin and eosin staining. Adipocyte size area was measured by outlining the adipocytes, and the average of five representative 10X fields from each mouse were plotted for cross section area distribution, as described (Xiong et al., 2021).

### Indirect calorimetry

Whole-body energy homeostasis was analyzed using the Promethion Core Metabolic System (Sable Systems) by the City of Hope Comprehensive Metabolic Phenotyping Core. Mice were single housed and acclimated in metabolic cages, with ad libitum food and water with controlled lighting for 2 days prior to metabolic recording. Metabolic parameters, including oxygen consumption, CO2 production, respiratory exchange ratio, ambulatory activity, and food intake were recorded for 5 consecutive days, as described (Yin et al., 2020).

### Statistical analysis

Data are presented as mean ± SE or SD as indicated. Each experiment was repeated at minimum twice to validate the result. Sample size were indicated for each experiment in figure legends. Statistical analysis was performed using GraphPad Prism. A minimum of three biological replicates were used to perform statistical analysis. Two-tailed Student’s t-test or One-way ANOVA with post-hoc analysis for multiple comparisons were performed as appropriate as indicated. P<0.05 was considered statistically significant.

## Results

### Nobiletin promotes clock oscillation in adipogenic progenitors

Nobiletin is known to increase clock oscillatory amplitude in skeletal muscle (He et al., 2016). To determine whether Nob modulates clock properties in adipocytes, we generated two adipogenic clock reporter progenitor cell lines, mesenchymal stem cell C3H10T1/2 and the preadipocyte 3T3-L1, with stable expression of a *Per2::dLuc* luciferase reporter. Using these reporter adipogenic progenitor cell lines, we examined the effect of Nob treatment on clock function via continuous monitoring of luciferase activity for 6 days. In 3T3-L1 reporter cells containing *Per2::dLuc*, Nob treatment at 1 to 5 μM induced a dose-dependent increase in amplitude (Fig. 1A & 1B), together with prolonged period length (Fig. 1C). Similar effects of Nob was observed in the C3H10T1/2 reporter line containing *Per2::dLuc* with enhanced cycling amplitude and increased period (Fig. 1D-1F), suggesting activation of clock oscillation. Consistent with its modulation of clock properties, Nob induced up-regulation of core clock genes in 3T3L1 adipogenic progenitors, including *Bmal1*, and negative clock regulators *Cry1, Per1* and *Per2* (Fig. 1G). Similar Nob inductions of clock genes in C3H10T1/2 mesenchymal precursor were also observed, although mostly only significantly elevated at 5 μM (Fig. 1H). Collectively, these findings indicate that Nob was sufficient to promote clock function in adipogenic progenitors.

**Figure 1.**
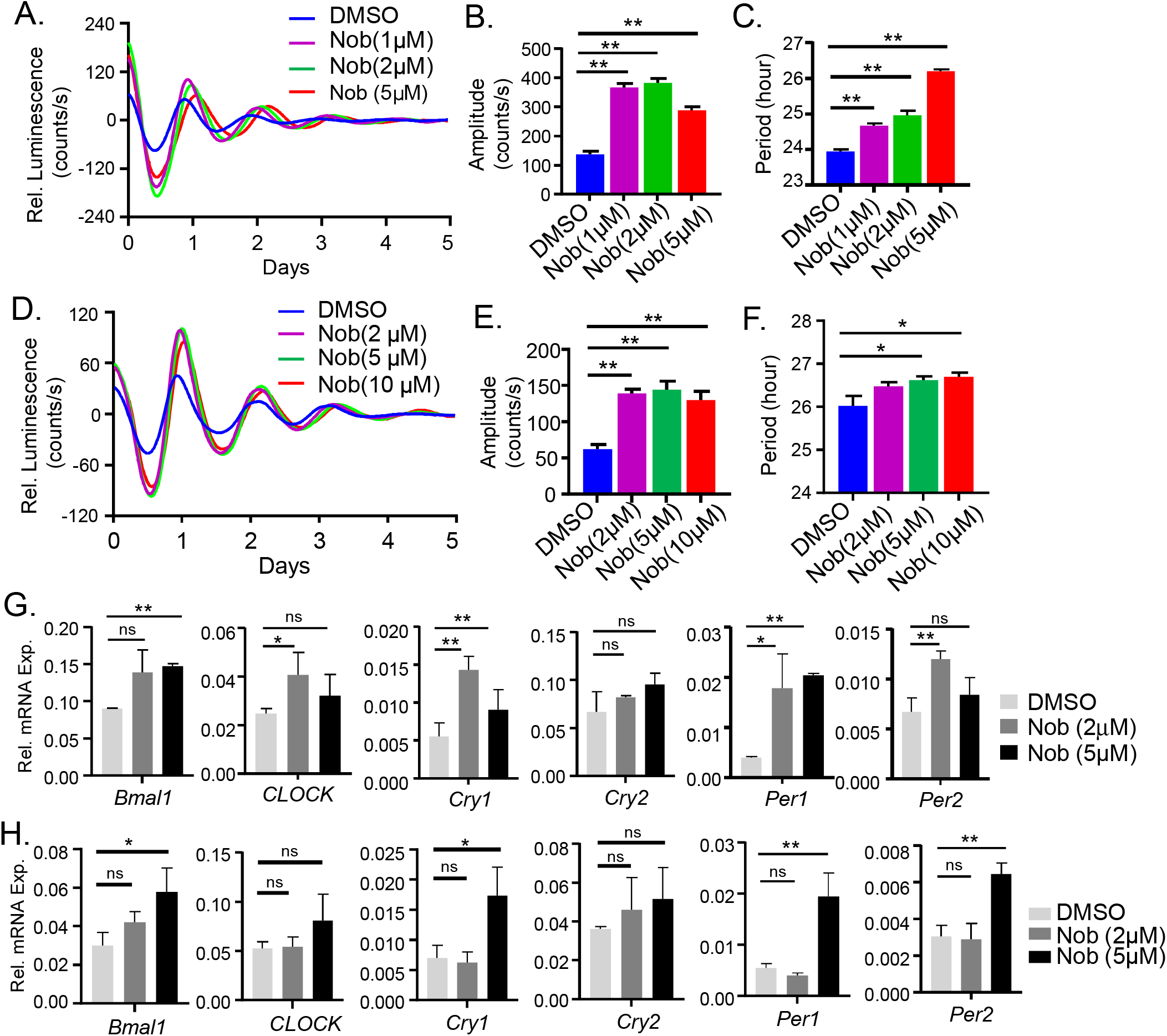
Nobiletin promotes clock oscillation in adipocytes. **(A-C)** Baseline-adjusted tracing plots of average bioluminescence activity of 3T3-L1 cells with stable expression of a *Per2::dLuc* reporter for 5 days (A), with quantitative analysis of clock cycling amplitude (B) and period length (C), with Nobiletin (Nob) treatment at indicated concentrations. (D-F) Baseline-adjusted tracing plots of average bioluminescence activity of C3H10T1/2 cells with stable expression of a *Per2::dLuc* reporter for 5 days (D), with quantitative analysis of clock cycling amplitude (E) and period length (F), with Nobiletin (Nob) treatment at indicated concentrations. Data are presented as Mean ± SD of n=4 replicates for each concentration tested, with three independent repeat experiments. (G, H) RT-qPCR analysis of clock gene expression at indicated concentrations of Nobiletin in 3T3-L1 cells (G) and C3H10T1/2 cells (H). Data are presented as Mean ± SD of n=3 replicates. *, **: p<0.05 and 0.01 Nob vs. DMSO by Student’s t test.

### Nobiletin inhibits the terminal differentiation and lineage commitment of adipogenic progenitors

We previously demonstrated that the positive clock regulator Bmal1 inhibits adipogenesis, whereas the clock repressor Rev-erba promotes this process (Guo et al., 2012; Nam et al., 2015a). The clock-modulatory properties of Nob in adipogenic progenitors led us to test its influence on their adipogenic potential. Indeed, Nob treatment of 3T3-L1 preadipocytes resulted in a dose-dependent reduction of lipid accumulation during adipogenic induction, as indicated by oil-red-O (Fig. 2A) and Bodipy staining (Fig. 2B). During adipogenic differentiation, the key adipogenic factors, C/EBPα and PPARγ, were robustly induced in control cells at day 6 as compared to before induction, as expected (Fig. 2C). In contrast, Nob markedly reduced their protein expression at day 6 of differentiation as compared to controls (Fig. 2C). Furthermore, Nob treatment attenuated the expression of mature adipocytes markers, fatty acid synthase (FASN) and the fatty acid transporter, fatty acid binding protein (FABP4), indicating impaired adipocyte terminal differentiation. Using the 10T1/2 adipogenic mesenchymal precursor cells, we further tested whether Nob modulates their lineage commitment and maturation to adipocytes. An analysis of differentiation via oil-red-O staining at early stage of day 5 (Fig. 3A) and mature differentiation at day 8 (Fig. 3B) showed that Nob suppressed the differentiation of immature mesenchymal precursor into mature adipocyte in a dose-dependent manner, with 5 μM displaying a stronger effect in preventing lipid accumulation during adipogenic progression. These findings were further validated by Bodipy staining demonstrating similar effects of Nob inhibition of adipogenic maturation (Fig. 3C). Consistent with these observations, an examination of adipogenic factors revealed robust reductions of C/EBPα and PPARγ protein levels by 5 μM Nob treatment (Fig. 3D), with similarly attenuated expression of FASN and FABP4 (Fig. 3E). Thus Nob displayed strong activity in inhibiting the development of adipogenic progenitor to mature adipocytes at distinct stages of lineage progression.

**Figure 2.**
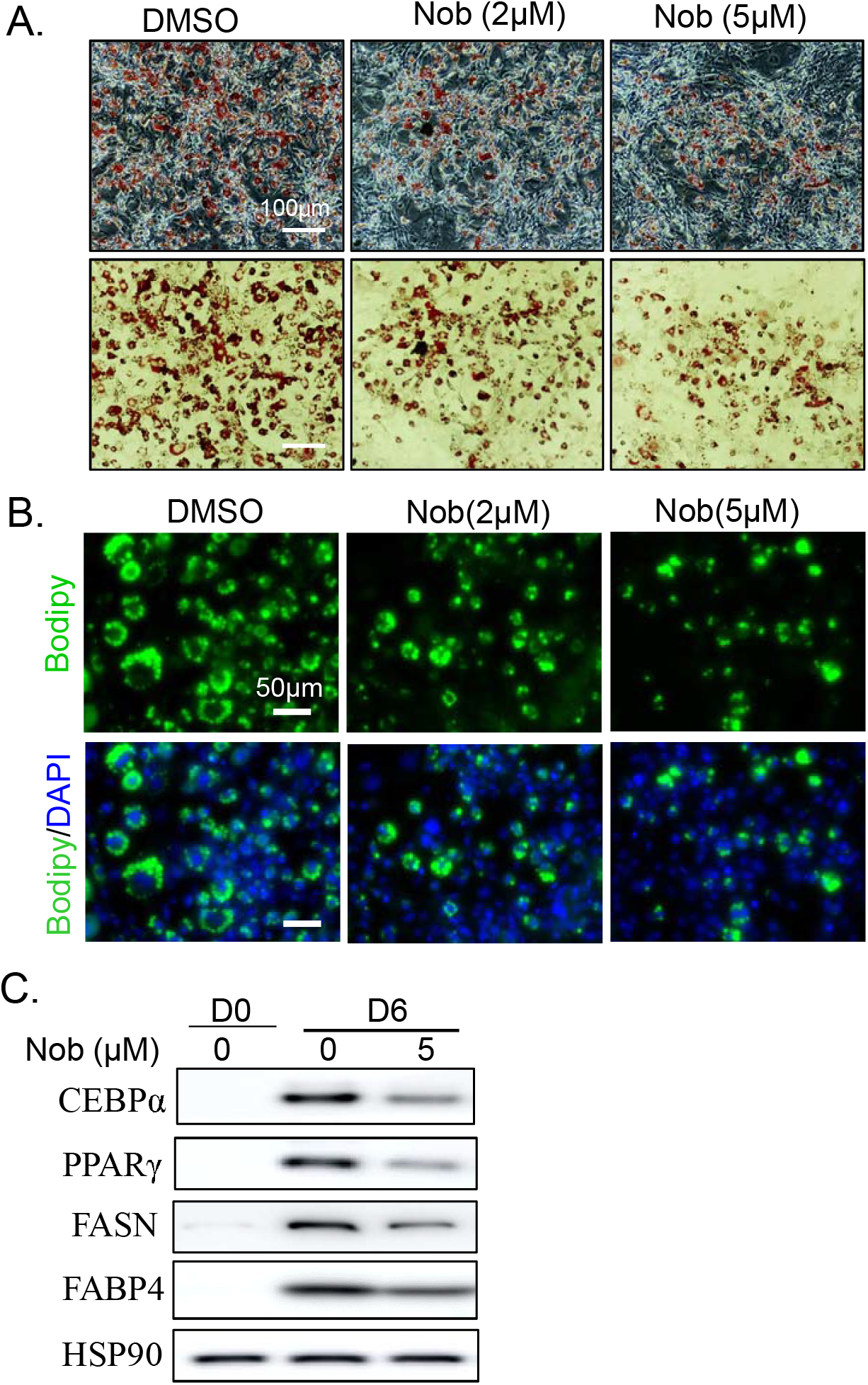
Effect of Nobiletin on terminal differentiation of 3T3-L1 preadipocyte. (A, B) Representative images of oil-red-O (A) and bodipy staining (B) of lipids at day 6 of 3T3-L1 preadipocyte differentiation with Nobiletin treatment at indicated concentrations. Scale bars: 100 μm (A) and 50 μm (B). ((C) Immunoblot analysis of adipogenic factors and mature adipocyte markers at Day 0 and day 6 of 3T3-L1 differentiation with Nobiletin treatment at indicated concentration. Each lane represents a pooled sample of 3 replicates.

**Figure 3.**
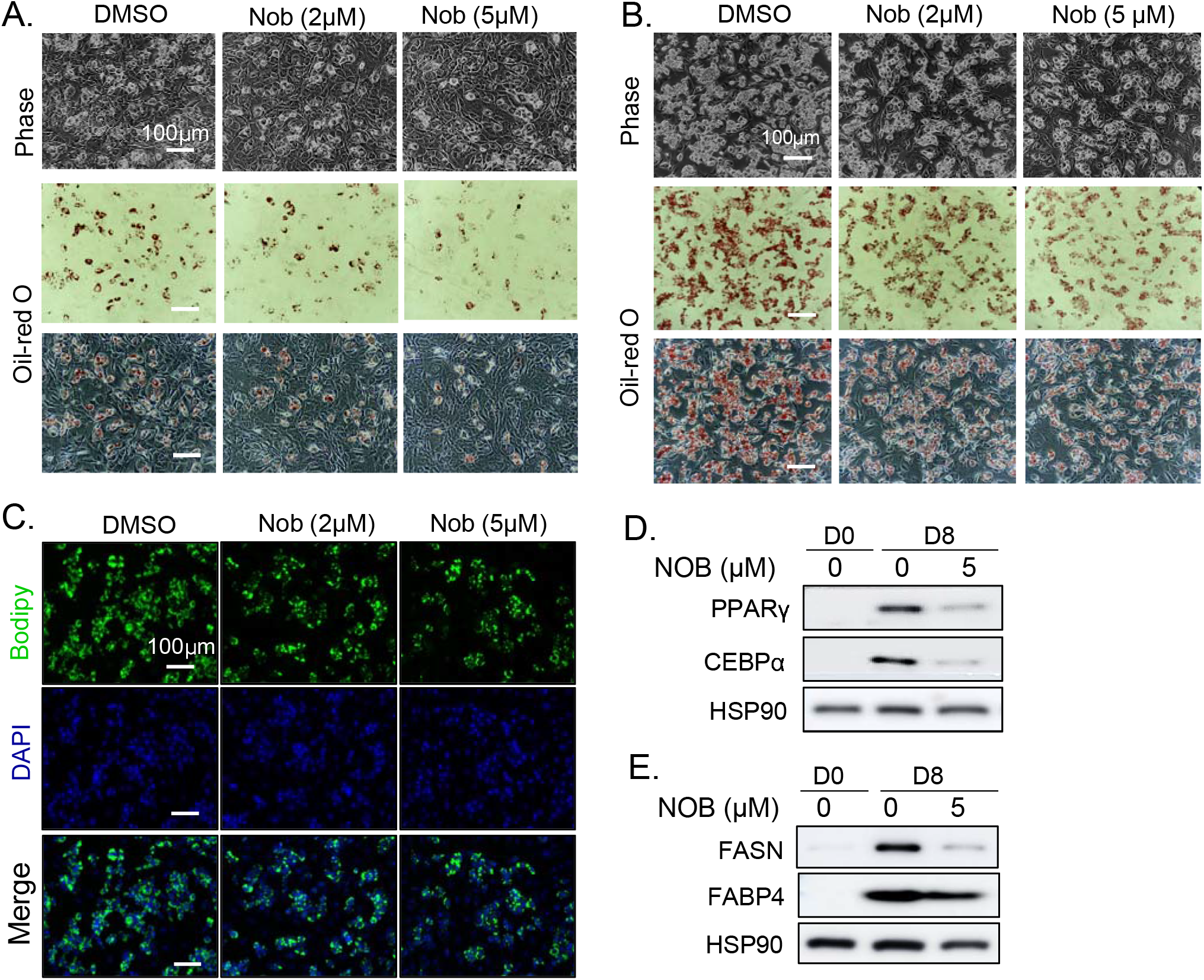
Effect of Nobiletin on adipogenic differentiation of C3H10T1/2 mesenchymal precursor cells. (A, B) Representative images of phase contrast and oil-red-O staining of lipid accumulation at day 5 (A) and day 8 (B) of C3H10T1/2 differentiation with Nobiletin treatment at indicated concentrations. Scale bars: 100 μm. (C) Representative images of Bodipy staining of lipids at day 8 of C3H10T1/2 differentiation with Nobiletin treatment at indicated concentrations. (D, E) Immunoblot analysis of adipogenic factors (D) and mature adipocyte markers (E) at Day 0 and day 8 of C3H10T1/2 differentiation with Nobiletin treatment at indicated concentration. Each lane represents a pooled sample of 3 replicates.

### Nobiletin induces key components of Wnt pathway and promotes Wnt signaling activity

The Wnt signaling pathway is a strong suppressor of adipocyte differentiation (Ref). Our previous studies demonstrated Bmal1 exerts a direct, clock-dependent transcriptional activation of Wnt signaling components to inhibit adipogenesis. Based on its clock-enhancing activity in adipogenic progenitors, Nobiletin may up-regulate signaling components of the Wnt pathway and attenuate adipocyte development. Consistent with its effect on inducing Bmal1 expression, in both 3T3-L1 preadipocytes (Fig. 4A) and 10T1/2 mesenchymal precursors (Fig. 4B), Nob treatment, at 2 and 5 μM, was sufficient to induce the mRNA levels of upstream Wnt signaling components, including the Wnt ligand *Wnt10a*, and Wnt receptors *Frizzed 1* or 2 (*Fzd1/2*) and *Disheveled 2* (*Dv12*) Furthermore, the Wnt signaling transcription effector, *Tcf4* was induced in 10T1/2 cells (Fig. 4B) but not 3T3-L1 preadipocytes (Fig. 4A), suggesting differential Nob regulation of Wnt pathway in distinct adipogenic cell types. Notably, the central signaling mediator of the Wnt activity, *β-catenin*, was up-regulated by Nob in both cell types, although interestingly this effect was only significant at 2 μM. β-catenin protein was suppressed during the adipogenic differentiation of control 10T1/2 cells from beginning of induction to day 8, reflecting the expected loss of Wnt activity during adipocyte development (Fig. 4C). In contrast, Nob treatment reversed this suppression of β-catenin expression at day 8 of 10T1/2 adipogenic differentiation. This suggested that a re-induction of Wnt signaling activity could underlie the inhibitory effect of Nob on adipogenesis. To test the direct effect of Nob on Wnt signaling activity in adipogenic progenitors, we used a transient transfection of the 10T1/2 cells with the TOPFlash luciferase reporter containing a Wnt-responsive TCF4-driven promoter. Under basal, non-Wnt stimulated condition, Nob induced Wnt signaling activity by 3-4~fold (Fig. 4D). Wnt-containing media stimulated the reporter activity by ~20 fold, as expected, which was further enhanced by 4-fold at 2 and 5 uM of Nob, indicating that Nob-mediated transcriptional activation of Wnt pathway components augmented the stimulation of Wnt signaling in adipogenic progenitors.

**Figure 4.**
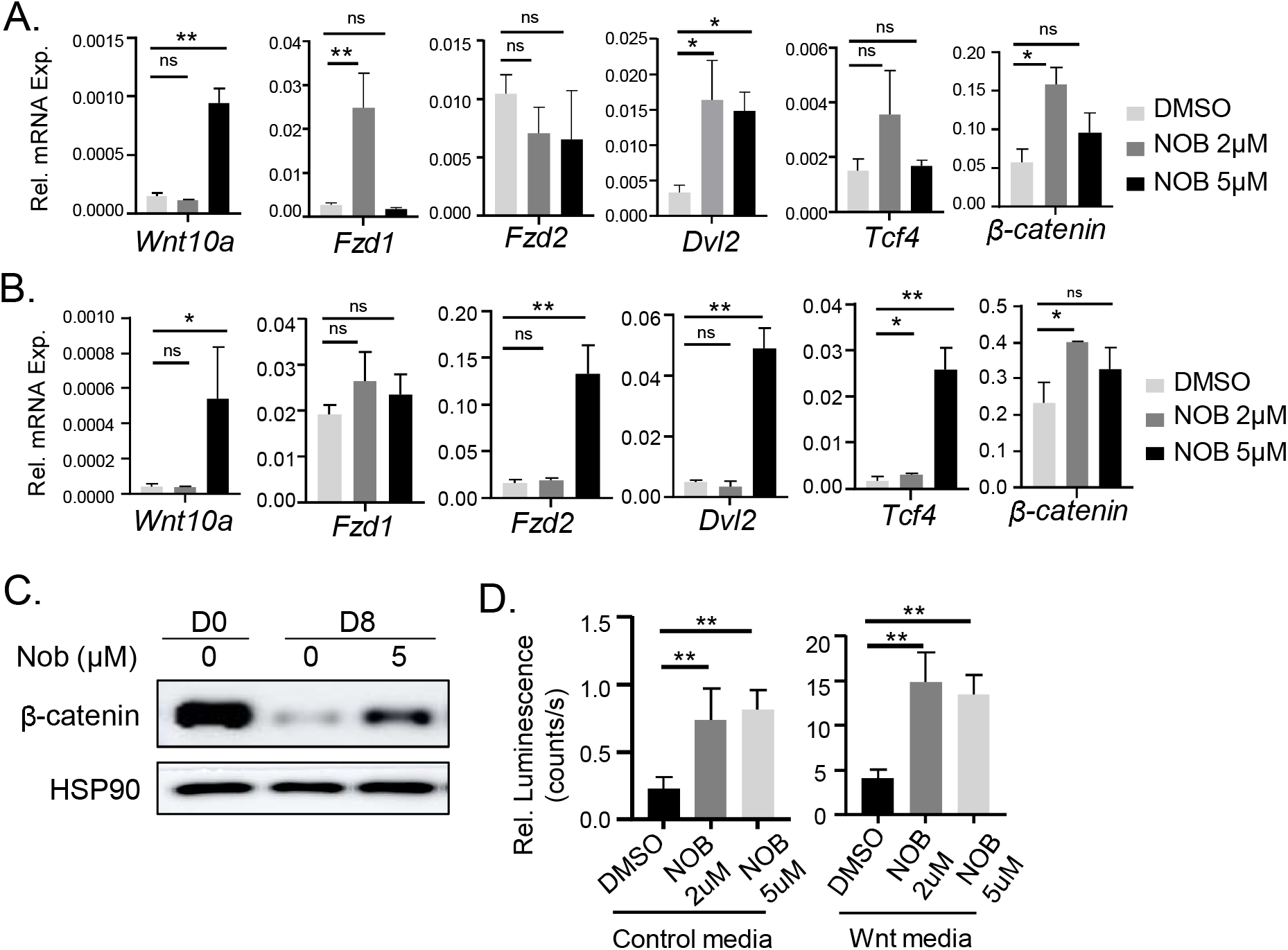
Nobiletin inhibits adipogenic differentiation via Wnt signaling pathway. (A, B) RT-qPCR analysis of expression of genes involved in the Wnt signaling pathway with Nobiletin treatment at indicated concentrations in 3T3-L1 preadipocytes (A) and C3H10T1/2 mesenchymal precursors (B). Data are presented as Mean ± SD of n=3 replicates. *, **: p<0.05 and 0.01 Nob vs. DMSO by Student’s t test. (C) Immunoblot analysis of β-catenin at day 0 and 8 of C3H10T1/2 adipogenic differentiation with Nobiletin treatment. Each lane represents a pooled sample of 3 replicates. (D) Analysis of Nobiletin effect on TOPFlash luciferase reporter activity in C3H10T1/2 the presence or absence of 10% Wnt3a media at indicated concentrations. Data are presented as Mean ± SD of n=4 replicates. **: p<0.05 and 0.01 Nob vs. DMSO by Student’s t test.

### Nobiletin treatment in mice reduces adiposity and adipocyte hypertrophy

Based on the suppression of Nob on adipogenic differentiation, we tested its potential effects on adipose tissue in vivo. Treatment of 48-week-old mice with custom 0.1% Nob-supplemented chow diet for 4 weeks led to a significant lowering of total body weight, compared to mice fed with a control chow diet (Fig. 5A). The reduced body weight in Nob-treated cohort was due to a lack of weight gain (Fig. 5B). The reduction of adipose tissue weight in these mice likely accounts for the lower body weight after 4 weeks of Nob diet (Fig. 5C), while muscle tissue weight was maintained (Fig. 5D). We next tested a 0.2% Nob-supplemented diet in younger 10-week-old mice and found a similar inhibition of the normal increase in body weight observed in the control cohorts during the 4-week treatment (Fig. 5E). Analysis of body composition by NMR revealed that the attenuated weight gain in the Nob-treated mice was likely attributed to loss of fat mass (Fig. 5F & 5G), which was significantly lower after 4 weeks compared to the control cohort. Total muscle mass was maintained for the Nob-treated mice with the ratio to body weight significantly higher, a consequence of reduction of fat content (Fig. 5H). Examination of whole-body energy balance during the last week of Nob treatment revealed comparable rates of food intake between control and Nob-treated mice (Fig. 5I & 5J), suggesting the loss of fat induced by Nob diet is not due to potential effect on food consumption. Interestingly, Nob treatment moderately reduced the oxygen consumption rate in mice under both light and dark cycles (Fig. 5K & 5L), with a similar trend of CO2 output (data not shown). In addition, a moderately increased locomotor activity confined to the light cycle was observed in the Nob-treated groups (Fig. 5M & 5N). We collected distinct fat depots for direct examination of Nob effects on adipocyte histology in the classic epidydimal visceral white adipose tissue (eWAT), subcutaneous inguinal white adipose tissue (iWAT) and the classic interscapular brown adipose tissue (BAT). A 0.2% Nob diet feeding for 4 weeks markedly reduced the amount of white fat pads from eWAT and iWAT (Fig. 6A & 6B), while this effect was not evident in BAT (Fig. 6C). The ratio of Tibialis Anterior (TA) muscle weight to total body weight was higher in the Nob diet-treated cohort than controls (Fig. 6D), possibly due to the loss of fat mass with a reduction of total body weight. Consistent with the significant reduction of adipose tissue mass, histological analysis revealed markedly smaller adipocytes in eWAT and iWAT fat pads by Nob treatment (Fig. 6E). Although BAT weight was not altered by Nob treatment, brown adipocytes displayed marked loss of lipids accumulation as indicated by H/E histology. Furthermore, quantification of adipocyte cell size distribution revealed a significant shift toward smaller adipocytes for both visceral eWAT and subcutaneous iWAT adipose depots (Fig. 6F & 6G), demonstrating that Nob treatment led to a marked reduction in adipocyte hypertrophy.

**Figure 5.**
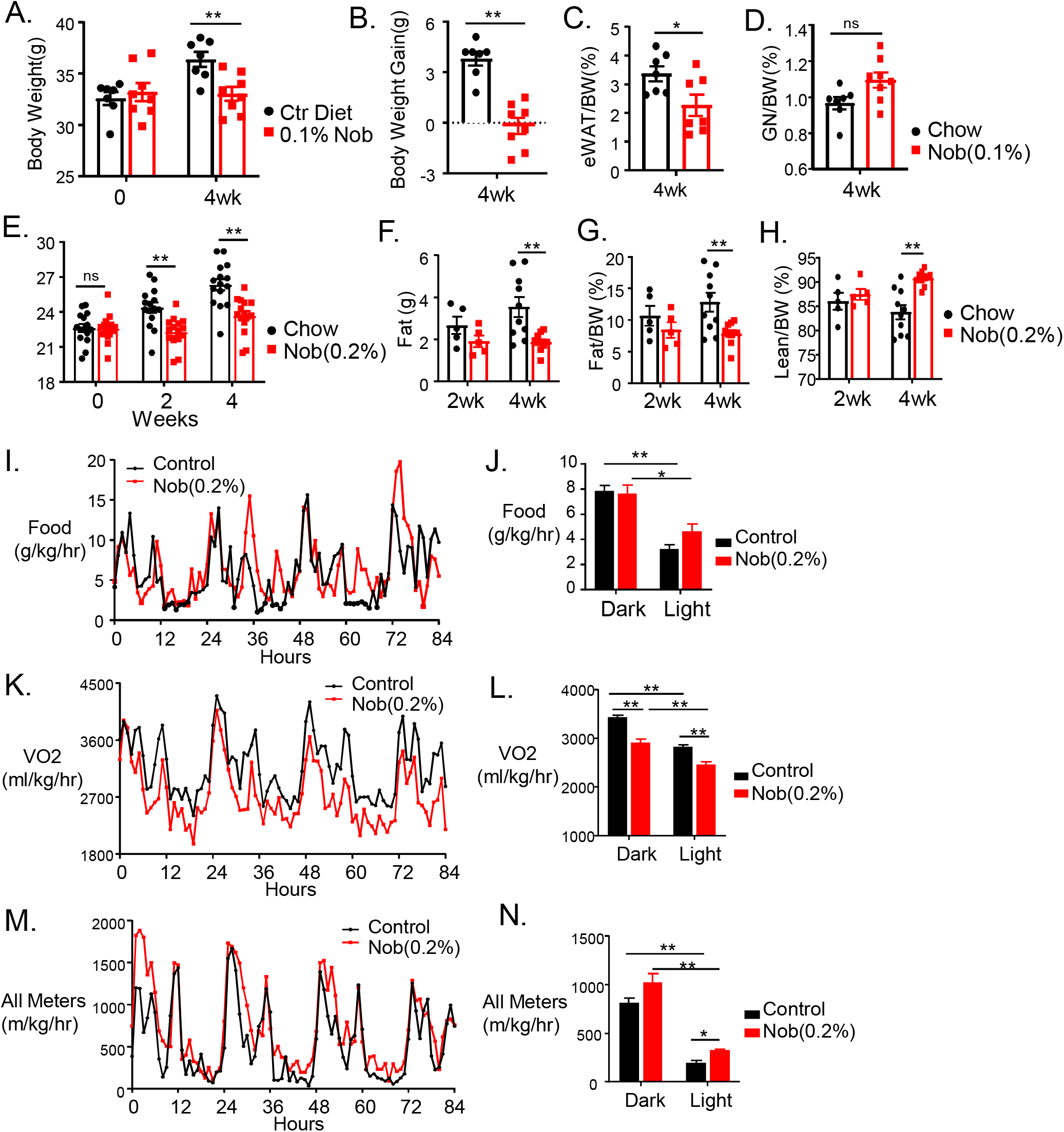
Nobiletin displays anti-obesity effect in vivo. (A-D) Analysis of the effect of 0.1% Nob-supplemented diet feeding for 4 weeks on body weight (A), change in body weight (B), epidydimal white adipose tissue (eWAT, C) and gastrocnemius (GN) muscle weight to body weight ratio (D) in 40-week-old male mice (n=7/group). (E-H) Effect of 0.2% Nob-supplemented diet feeding for 4 weeks on body weight (E), and NMR analysis of total fat mass (F), total fat mass to body weight ratio (G) and lean mass to body weight ratio (H) in 10-week-old male mice (n=5-10/group). (I-N) Analysis of the effect of 0.2% Nob-supplemented diet feeding on whole-body energy homeostasis. Average plots with quantitative analysis of food intake (I, J), oxygen consumption (K, L) and total activity monitoring (M, N) in control and Nobiletin-treated mice. n=5 mice/group. Data are presented as Mean ± SE. *, **: p<0.05 and 0.01 Nob vs. control diet by Student’s t test.

**Figure 6.**
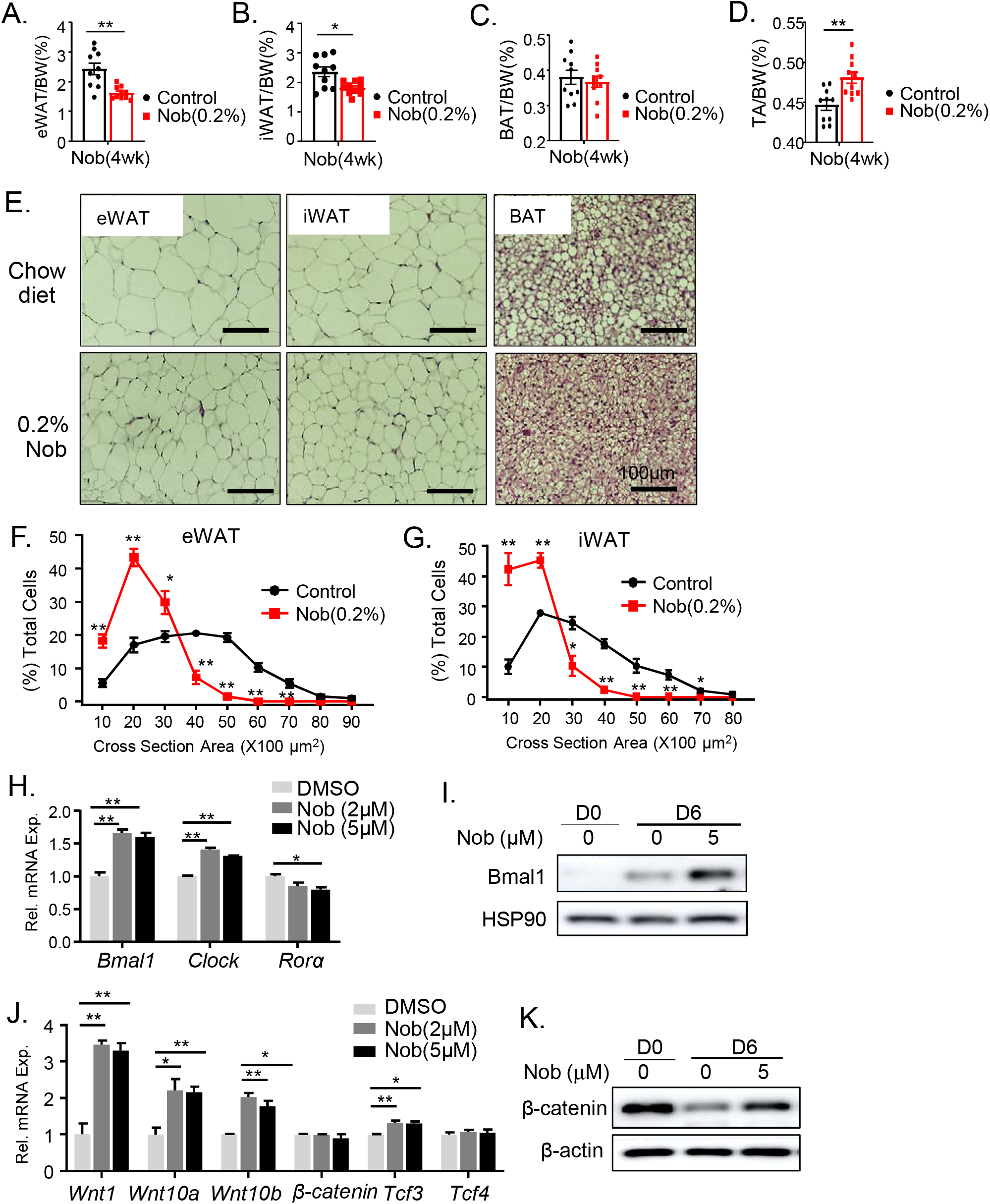
Nobiletin administration in mice reduces adipose tissue expansion. (A-D) Analysis of effect of 0.2% Nob-supplemented diet feeding on tissue weight after 4 weeks of treatment in epidydimal white adipose tissue (eWAT, A), inguinal subcutaneous white adipose tissue (iWAT, B), interscapular brown adipose tissue (iBAT, C), and Tibialis Anterior (TA, D). (n=10/group). (E) Representative H/E histology images of eWAT, iWAT and BAT in chow and 0.2% Nob-supplemented diet mice after 4 weeks. Scale bar: 100 μm. (F, G) Analysis of adipocyte cell size distribution of eWAT (F) and iWAT (G) in normal chow control and 0.2% Nob-supplemented diet mice. (H) RT-qPCR analysis of circadian clock genes induction by Nob in isolate primary preadipocytes at indicated concentrations. Data are presented as Mean ± SD of n=3 replicates. *, **: p<0.05 and 0.01 Nob vs. DMSO by Student’s t test. (I) Immunoblot analysis of Bmal1 protein expression by 5μM Nob treatment at day 0 and 6 of adipogenic induction in primary preadipocytes. (J) RT-qPCR analysis of Wnt pathway genes by Nob in isolate primary preadipocytes at indicated concentrations. Data are presented as Mean ± SD of 3 replicates. *, **: p<0.05 and 0.01 Nob vs. DMSO by Student’s t test. (K) Immunoblot analysis of β-catenin protein expression by 5μM Nob treatment at day 0 and 6 of adipogenic induction of primary preadipocytes. Each lane represents a pooled sample of 3 replicates.

To determine the mechanism underlying the *in vivo* effects of Nob on preventing adipose tissue expansion, we isolated the stromal vascular fraction containing adipogenic progenitors from iWAT depots and determined their differentiation potential in response to Nob. Nob was sufficient to induce *Bmal1* and *Clock* gene expression (Fig. 6H), with robust increased in Bmal1 protein in response to 5μM Nob when the primary preadipocytes were subjected to adipogenic differentiation for 6 days (Fig. 6I). Interestingly, Nob had a moderate effect on repressing RORα, suggesting a potential negative feedback due to its agonist activity (Fig. 6H). In line with the findings of Nob modulation of Wnt pathway genes in adipogenic progenitor cell lines, it similarly induced key components of this cascade in primary preadipocytes. These included the Wnt ligands (*Wnt1, Wnt10a* and *Wnt10*), while *β-catenin, Tcf4*, and the Wnt receptor genes were not significantly altered (Fig. 6J). Consistent with this transcriptional regulation of Wnt pathway, Nob treatment restored the reduced β-catenin protein expression during preadipocyte differentiation, suggesting enhanced Wnt signaling activity induced by Nob (Fig. 6K).

### Nobiletin activates Wntpathway and inhibits adipogenic maturation via clock modulation

Our findings from distinct adipogenic progenitor cells lines indicated that Nob is sufficient to re-activate Wnt pathway during adipogenic differentiation. We thus postulated that the activation of Wnt signaling by Nob may inhibit the differentiation of primary preadipocytes into mature adipocyte. Indeed, 5 μM Nob robustly inhibited the formation of mature adipocytes, as indicated by the oil-red-O staining analysis of differentiation of primary preadipocytes at day 4 and day 6, (Fig. 7A & 7B). Immunoblot analysis of adipogenic factor expression indicated the downregulation of C/EBPα and C/EBPα along with an induction of Bmal1, while PPARγ was not altered (Fig. 7C). Consistent with the attenuated adipogenic maturation as assessed by lipid accumulation, Nob treatment resulted in near complete loss of mature adipocyte markers FASN and a strong inhibition of FABP4 (Fig. 7D). Lastly, to test whether Nob effect on inhibiting adipogenesis is mediated via its clock-modulatory activity, we used primary preadipocytes from iWAT-selective *Bmal1-null* (BMKO) along with controls from Floxed mice (BMCtr) to specifically address this (Xiong et al., 2022a). Bodipy staining showed that, as expected, Nob treatment efficiently suppressed the adipogenic differentiation of preadipocytes from control mice (Fig. 7E). In comparison, this inhibitory effect of Nob on adipocyte maturation was largely abolished in the *Bmal1-null* preadipocytes, indicating that Nob activity in suppressing adipogenesis is dependent on a Bmal1-driven functional clock.

**Figure 7.**
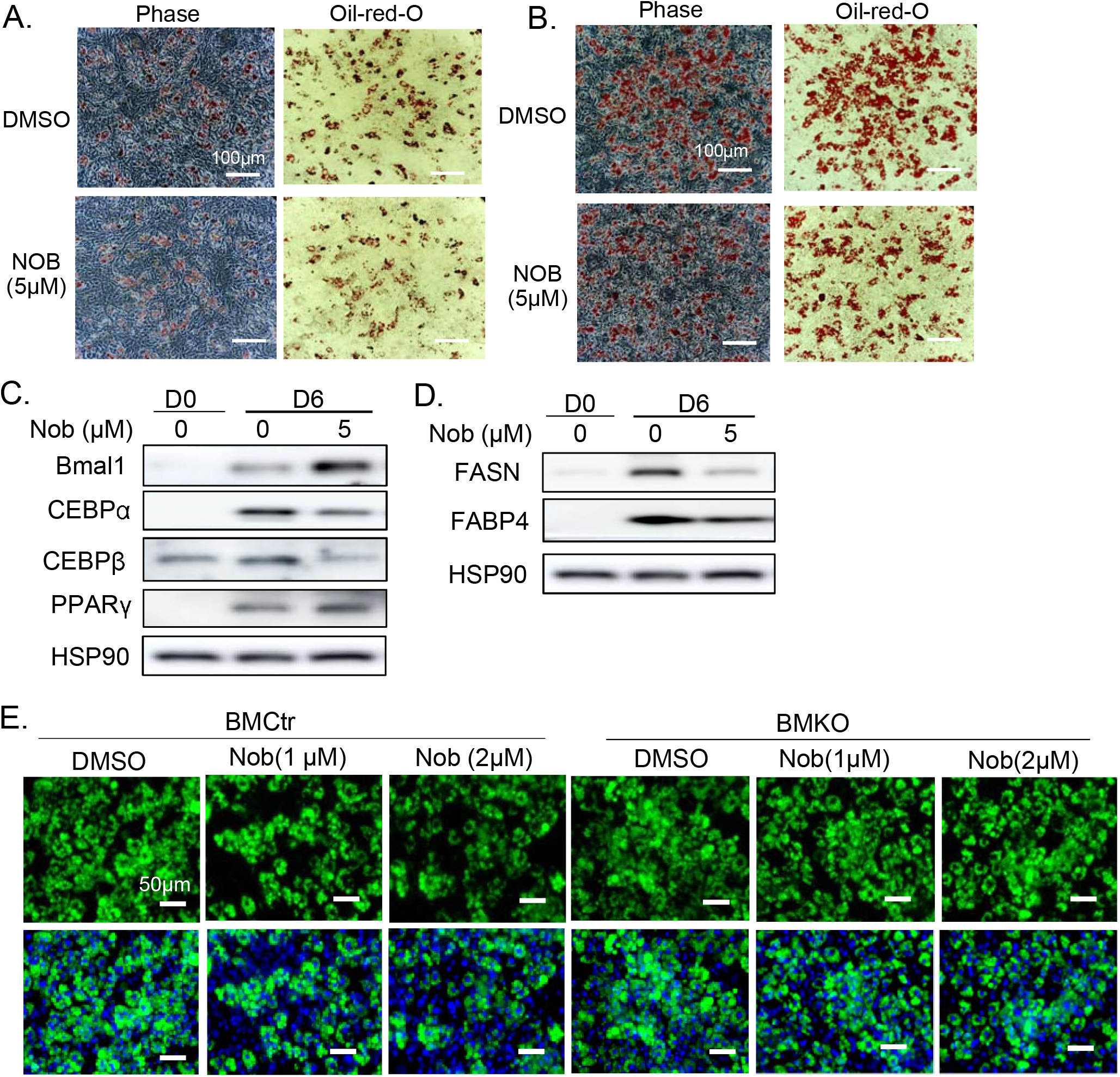
Effect of Nobiletin on primary preadipocyte differentiation. (A, B) Representative images of oil-red-O staining of adipogenic differentiation of isolated primary preadipocyte from the stromal vascular fraction of subcutaneous inguinal adipose tissue treated at indicated concentration of Nobiletin at day 4 (A) and day 6 (B). Scale bar: 100 μm. (C, D) Immunoblot analysis of adipogenic factors (C) and mature adipocyte markers (D) at day 0 and 6 of primary preadipocyte differentiation with Nobiletin treatment. Each lane represents a pooled sample of 3 replicates. (E) Representative images of Bodipy staining at day 6 of differentiation of primary preadipocytes isolated from floxed control (BMCtr) and Bmal1-deficient (BMKO) mice treated at indicated concentration of Nobiletin. Scale bar: 50 μm.

## Discussion

Despite previous reports of the metabolic activities of Nobiletin, its direct involvement in adipogenesis has not been investigated. Employing distinct adipogenic progenitor models, our study revealed a previously unappreciated activity of Nob in suppressing adipocyte formation that may contribute to its *in vivo* anti-obesity effect. We further demonstrate that the Nob inhibition of adipogenesis occurs through a Bmal1-dependent clock control of the Wnt signaling pathway.

Nob has been reported to display numerous metabolic benefits, with a wealth of current literature on its mechanism of action. Several previous studies noted an anti-obesity effect of Nob (He et al., 2016; Lee et al., 2010; Mulvihill et al., 2011), although the action of Nob in directly modulating adipocyte function has not been explored. Our findings add to the current understanding of Nob metabolic activities that involve its inhibitory effect on adipocyte differentiation. Due to its agonist activity for RORα/γ, Nob activates Bmal1 transcription via a direct RORE response element in its promoter (Preitner et al., 2002). RORα/γ, together with the transcription repressor Rev-erbaα, exert positive and negative modulations of RORE-mediated Bmal1 transcription, respectively, to elicit the oscillatory rhythm of this key clock transcriptional activator. Our analysis of clock modulatory activities of Nob in two distinct adipocyte progenitor cell types demonstrate that exert it is sufficient to enhance cell-autonomous adipocyte clock oscillation, consistent with its function as a RORα/γ agonist that activates Bmal1 transcription. Notably, in addition to its amplitude-enhancing effect as previously reported, our analysis in these cells revealed a consistent effect of Nob on period lengthening. Examination of clock gene modulation uncovered induction of key components of the negative feedback arm of core clock feedback loop, Per1/2 and Cry1/2, which may account for the increase in period length. Whether the effect of Nob on these clock genes is mediated by direct ROR transcriptional control or secondary to Bmal1 up-regulation remains to be dissected.

Our previous studies demonstrated a function of the circadian clock in modulating adipogenesis(Nam et al., 2015a; Nam et al., 2015b; Nam et al., 2016; Xiong et al., 2021) Transcriptional activator of the molecular clock circuit, Bmal1, inhibits adipogenesis (Guo et al., 2012). In the adipogenic differentiation models we tested, Nob induced Bmal1 transcription due to ROR activation. As Nob treatment led to up-regulation of Wnt pathway components that are direct Bmal1 targets, this effect could likely occur via Bmal1induction of. The observed inhibitory activity of Nob on adipogenesis supports this notion, and the loss of this effect in Bmal1-null preadipocytes further demonstrates the involvement of Bmal1 modulation. RORα has been shown to interact with C/EBPβ that blocks adipogenesis (Ohoka et al., 2009). It is conceivable that Nob, as an agonist of RORα, may function through this mechanism to enhance RORα interaction with C/EBPβ that suppresses adipogenic differentiation. This mechanism may act synergistically with the modulation of Bmal1-controlled Wnt signaling. How these disparate mechanisms contribute to the anti-adipogenic actions of Nob could be further dissected in future studies.

The strong effect of Nob in inducing fat loss in mice that led to reduction of total body weight remains intriguing. Although our findings suggest that inhibition of adipocyte formation may contribute, at least in part, to this *in vivo* action, additional mechanisms could apply. Analysis of energy balance in Nob-treated cohorts revealed no significant impact on food consumption, ruling out a potential central nervous system influence on appetite. However, Nob reduced oxygen consumption with moderately elevated activity level that was confined to the light cycle. The overall changes involving whole-body energy homeostasis observed here suggest that altered energy balance may not account for Nob effect on preventing weight gain in mice.

The direct action of Nob in inhibiting adipocyte development and its anti-obesity effects suggest that this natural compound may have potential therapeutic applications for managing obesity and prevention of the associated metabolic complications. The effect of Nob on augmenting clock oscillation could be beneficial for ameliorating shiftwork-associated obesity with potential public health impact. Towards drug development for new anti-obesity molecules, Nob may be leveraged as a lead compound for new derivatives with anti-obesity efficacy. As a ROR agonist that modulates clock oscillation with disparate properties attributing to to its metabolic benefits, dissecting Nob modulation of metabolic pathways that are clock-dependent or independent may provide mechanistic insights for rational anti-obesity drug design. Circadian disruption is strongly linked with the development of metabolic disorders (Pan et al., 2011), particularly obesity and insulin resistance that predispose to diabetes (Sinturel et al., 2020). Shiftwork increases the risk for Type II diabetes, and mice subjected to shiftwork regimen developed pronounced obesity with marked insulin resistance (Xiong et al., 2021). The prevalence of circadian misalignment in a modern lifestyle may contribute to the current epidemic of metabolic disorders (Pan et al., 2011; Roenneberg et al., 2012). With the elucidation of the inhibitory effect on adipocyte development and its clock-modulatory activity, Nob could be leveraged as a pharmacological clock intervention to prevent metabolic consequences associated with circadian misalignment.

## Acknowledgements

We thank Drs. Steve Kay and Meng Qu at the University of Southern California for sharing the luciferase reporters used in this study. We also thank the City of Hope Shared Resources Core Facility in carrying out metabolic phenotyping analysis. KM is a faculty member supported by the NCI-designated Comprehensive Cancer Center at the City of Hope National Cancer Center. This project was supported by a grant from National Institute of Health 1R01DK112794 to KM. and R01DK097160 and R01DK128972 to VY. The funders had no role in study design, data collection and analysis, decision to publish, or preparation of the manuscript.

## Authorship Statement

XX, TK and WL: data curation and investigation, formal analysis, manuscript editing; JL and VY: data curation and manuscript editing; KM: formal analysis, project administration, manuscript writing and editing, and funding acquisition.

