## Supplemental Figures for "The clock-modulatory activity of Nobiletin suppresses adipogenesis via Wnt signaling"

### Slide 1
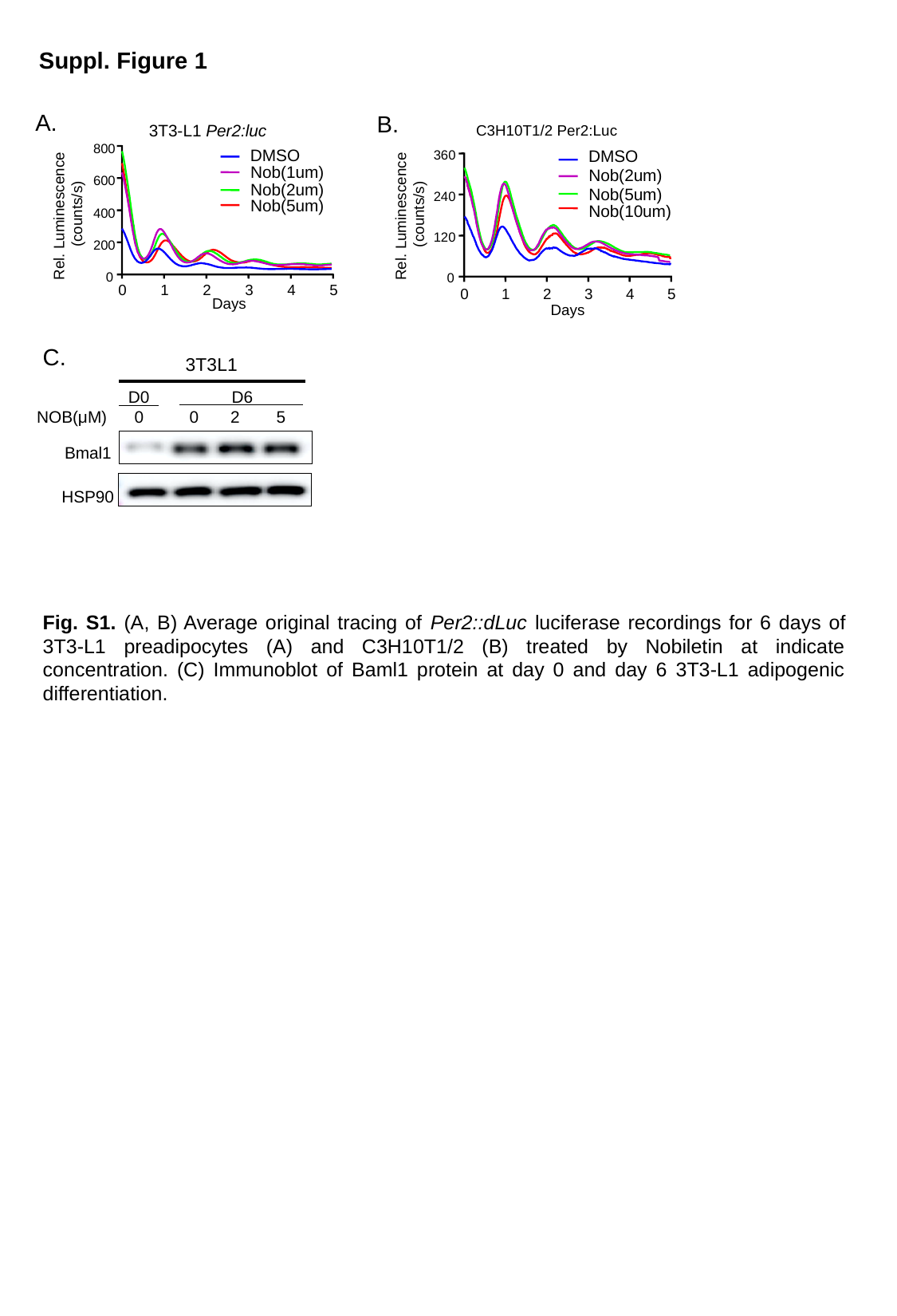

Suppl. Figure 1
A.
B.
3T3-L1 Per2:luc
800
600
400
200
0
0
1
2
3
4
5
Days
Rel. Luminescence
(counts/s)
DMSO
Nob(1um)
Nob(2um)
Nob(5um)
C3H10T1/2 Per2:Luc
DMSO
Nob(2um)
Nob(5um)
Nob(10um)
360
240
120
0
0
1
2
3
4
5
Days
Rel. Luminescence
(counts/s)
C.
 3T3L1
 D0 D6
NOB(μM) 0 0 2 5
Bmal1
HSP90
Fig. S1. (A, B) Average original tracing of Per2::dLuc luciferase recordings for 6 days of 3T3-L1 preadipocytes (A) and C3H10T1/2 (B) treated by Nobiletin at indicate concentration. (C) Immunoblot of Baml1 protein at day 0 and day 6 3T3-L1 adipogenic differentiation.

### Slide 2
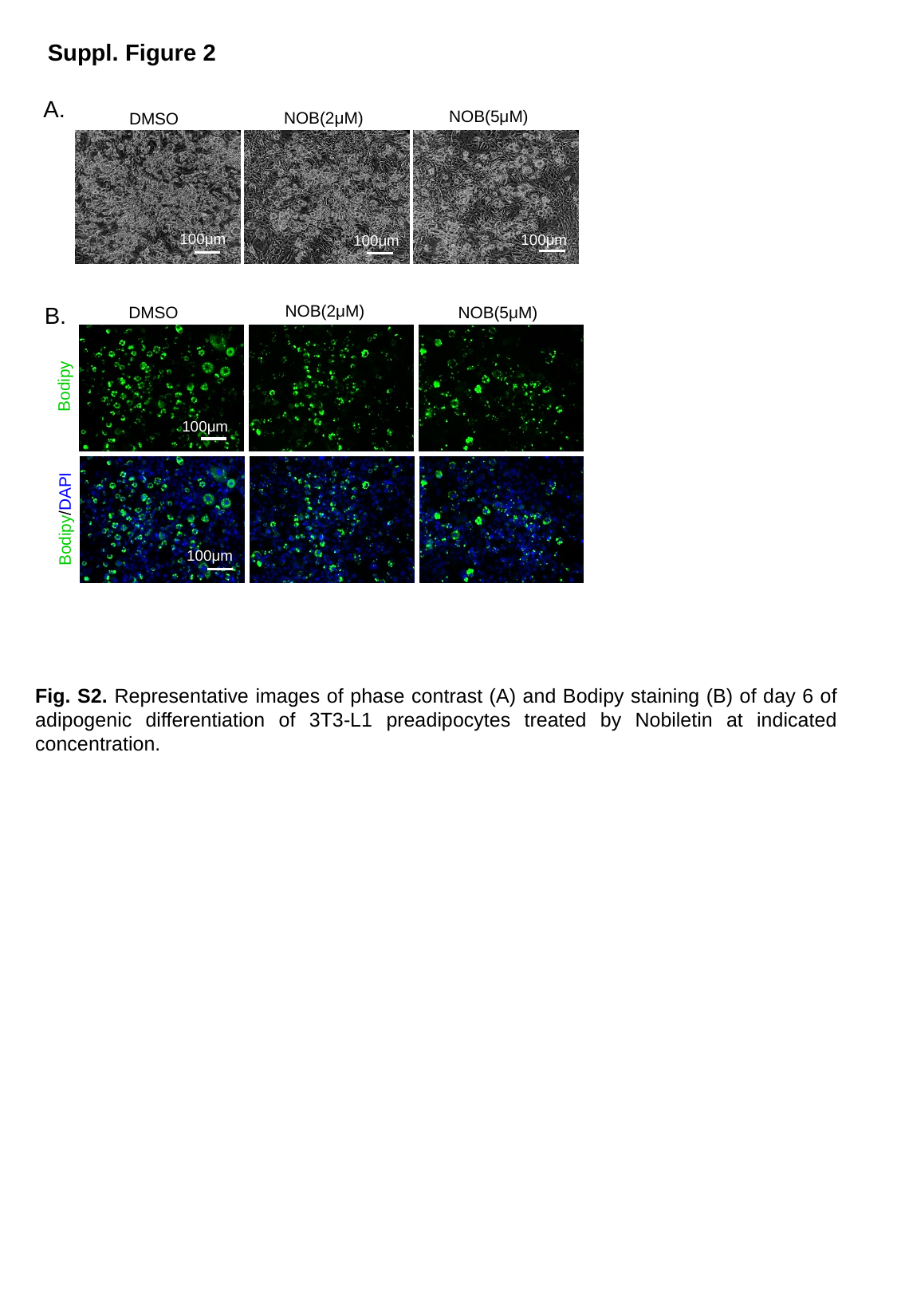

Suppl. Figure 2
A.
NOB(5μM)
NOB(2μM)
DMSO
100μm
100μm
100μm
NOB(2μM)
NOB(5μM)
DMSO
Bodipy
100μm
Bodipy/DAPI
100μm
B.
Fig. S2. Representative images of phase contrast (A) and Bodipy staining (B) of day 6 of adipogenic differentiation of 3T3-L1 preadipocytes treated by Nobiletin at indicated concentration.

### Slide 3
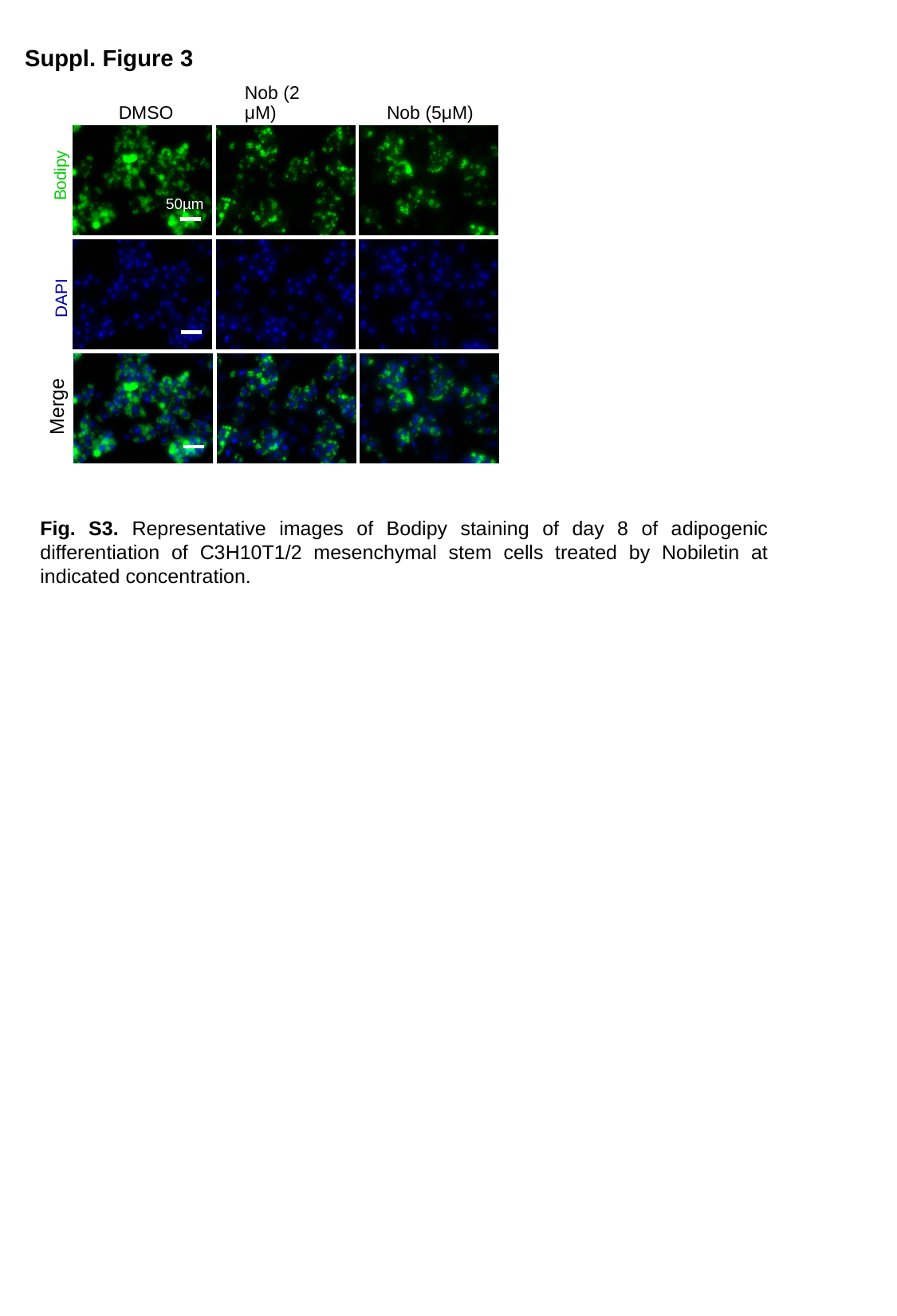

Suppl. Figure 3
DMSO
Nob (2 μM)
Nob (5μM)
Bodipy
50µm
DAPI
Merge
Fig. S3. Representative images of Bodipy staining of day 8 of adipogenic differentiation of C3H10T1/2 mesenchymal stem cells treated by Nobiletin at indicated concentration.

### Slide 4
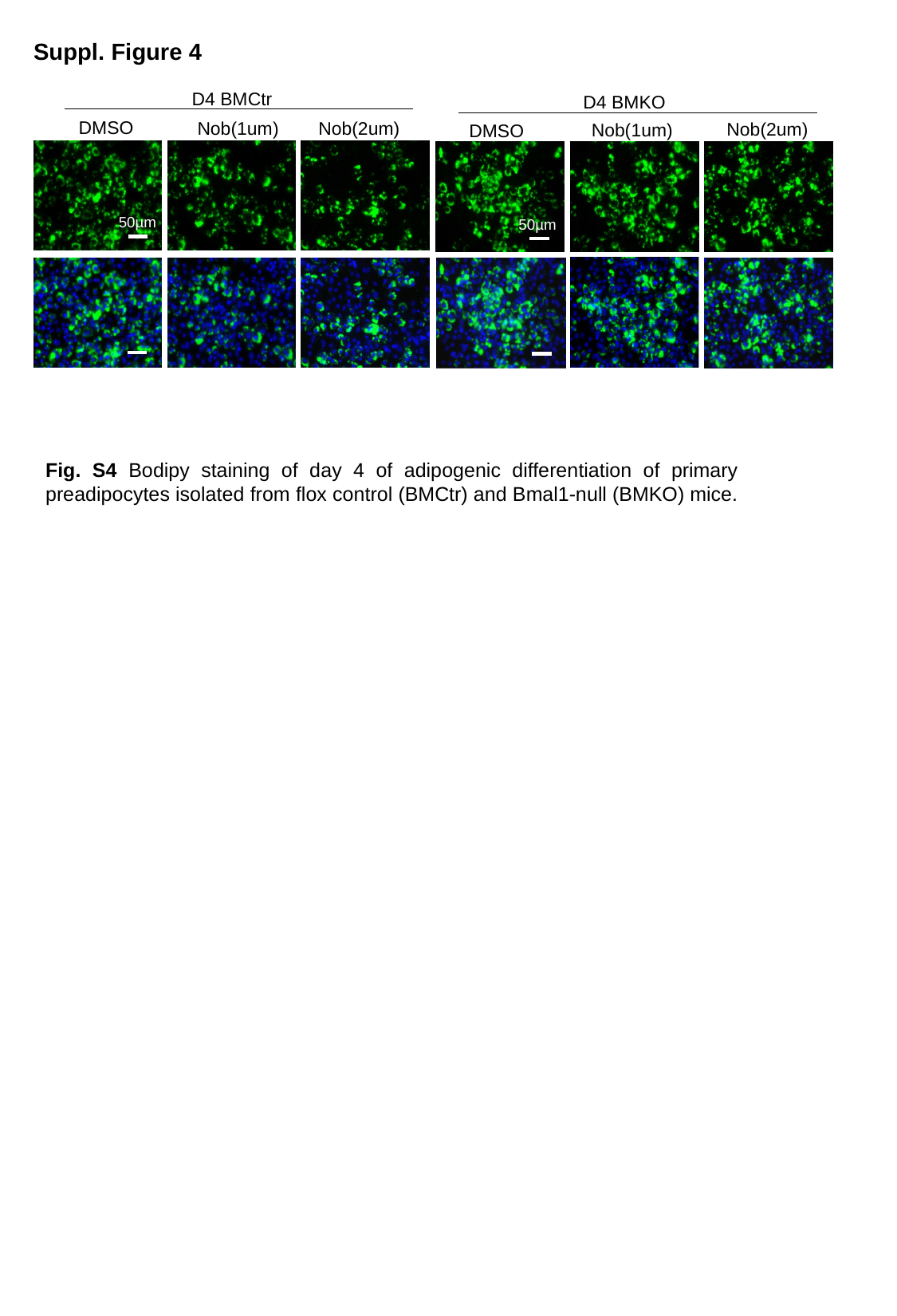

Suppl. Figure 4
D4 BMCtr
Nob(2um)
Nob(1um)
DMSO
50µm
50µm
D4 BMKO
50µm
Nob(1um)
Nob(2um)
DMSO
Fig. S4 Bodipy staining of day 4 of adipogenic differentiation of primary preadipocytes isolated from flox control (BMCtr) and Bmal1-null (BMKO) mice.

### Slide 5
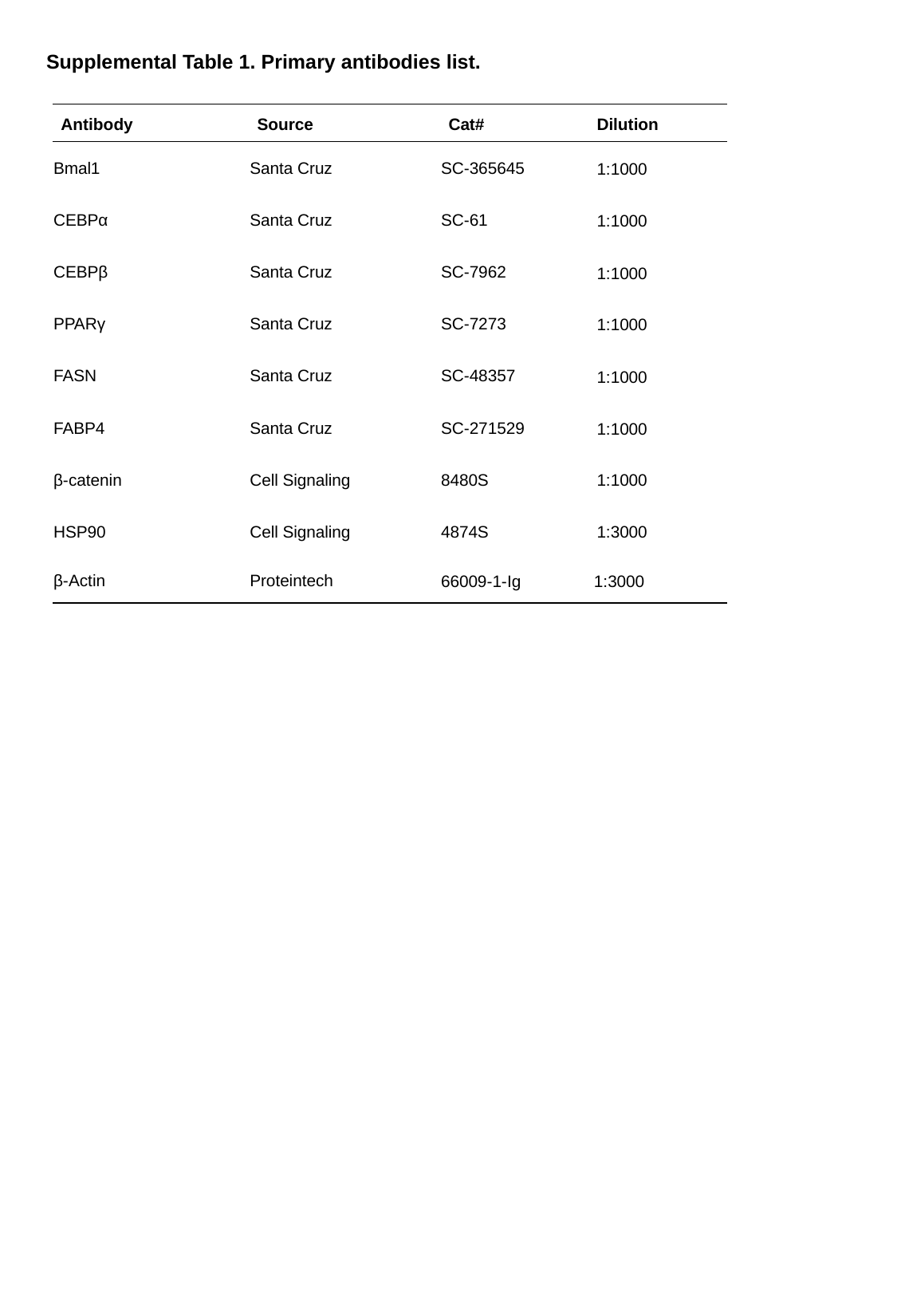

Supplemental Table 1. Primary antibodies list.
| Antibody | Source | Cat# | Dilution |
| --- | --- | --- | --- |
| Bmal1 | Santa Cruz | SC-365645 | 1:1000 |
| CEBPα | Santa Cruz | SC-61 | 1:1000 |
| CEBPβ | Santa Cruz | SC-7962 | 1:1000 |
| PPARγ | Santa Cruz | SC-7273 | 1:1000 |
| FASN | Santa Cruz | SC-48357 | 1:1000 |
| FABP4 | Santa Cruz | SC-271529 | 1:1000 |
| β-catenin | Cell Signaling | 8480S | 1:1000 |
| HSP90 | Cell Signaling | 4874S | 1:3000 |
| β-Actin | Proteintech | 66009-1-Ig | 1:3000 |

### Slide 6
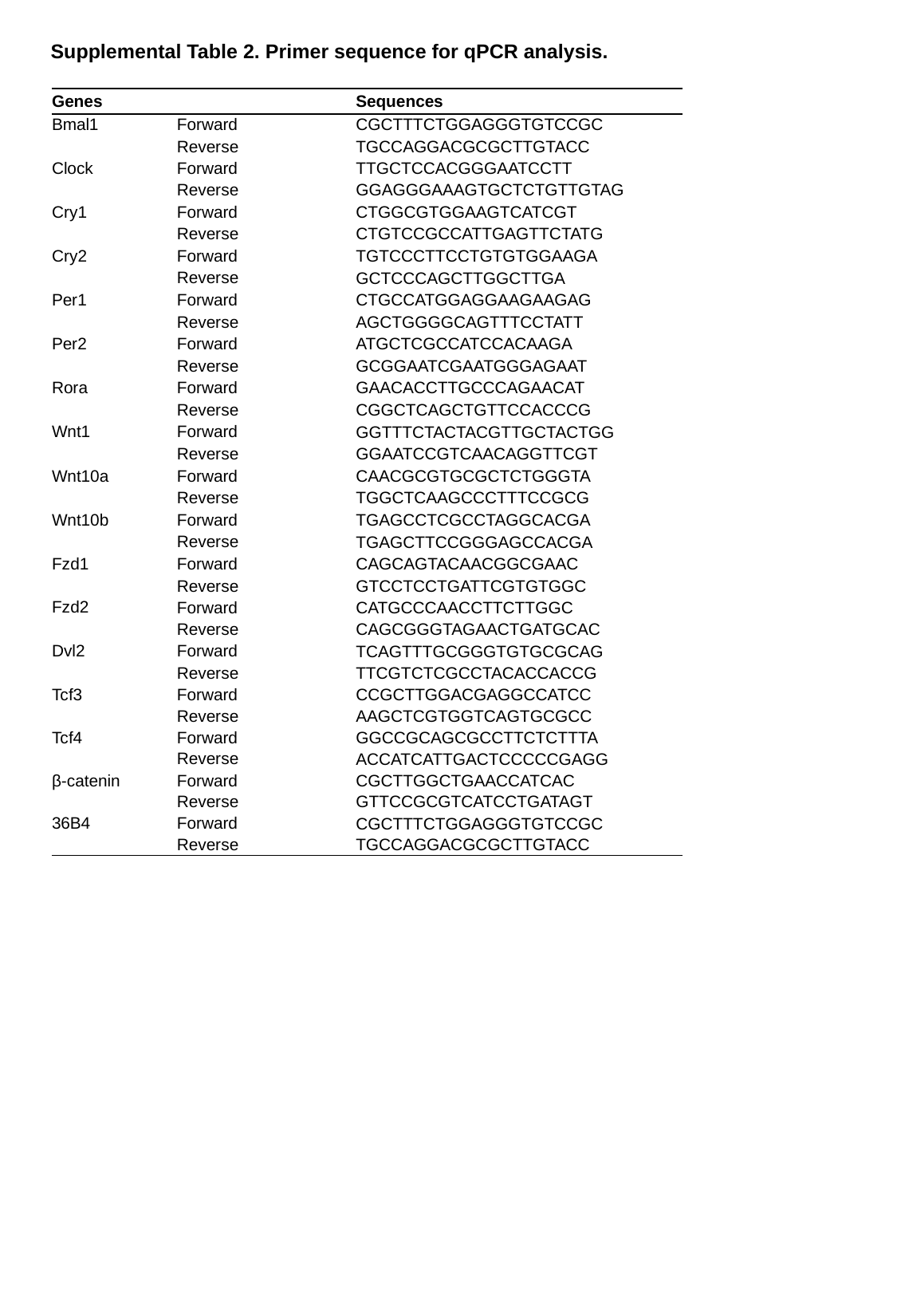

Supplemental Table 2. Primer sequence for qPCR analysis.
| Genes | | Sequences |
| --- | --- | --- |
| Bmal1 | Forward | CGCTTTCTGGAGGGTGTCCGC |
| | Reverse | TGCCAGGACGCGCTTGTACC |
| Clock | Forward | TTGCTCCACGGGAATCCTT |
| | Reverse | GGAGGGAAAGTGCTCTGTTGTAG |
| Cry1 | Forward | CTGGCGTGGAAGTCATCGT |
| | Reverse | CTGTCCGCCATTGAGTTCTATG |
| Cry2 | Forward | TGTCCCTTCCTGTGTGGAAGA |
| | Reverse | GCTCCCAGCTTGGCTTGA |
| Per1 | Forward | CTGCCATGGAGGAAGAAGAG |
| | Reverse | AGCTGGGGCAGTTTCCTATT |
| Per2 | Forward | ATGCTCGCCATCCACAAGA |
| | Reverse | GCGGAATCGAATGGGAGAAT |
| Rora | Forward | GAACACCTTGCCCAGAACAT |
| | Reverse | CGGCTCAGCTGTTCCACCCG |
| Wnt1 | Forward | GGTTTCTACTACGTTGCTACTGG |
| | Reverse | GGAATCCGTCAACAGGTTCGT |
| Wnt10a | Forward | CAACGCGTGCGCTCTGGGTA |
| | Reverse | TGGCTCAAGCCCTTTCCGCG |
| Wnt10b | Forward | TGAGCCTCGCCTAGGCACGA |
| | Reverse | TGAGCTTCCGGGAGCCACGA |
| Fzd1 | Forward | CAGCAGTACAACGGCGAAC |
| | Reverse | GTCCTCCTGATTCGTGTGGC |
| Fzd2 | Forward | CATGCCCAACCTTCTTGGC |
| | Reverse | CAGCGGGTAGAACTGATGCAC |
| Dvl2 | Forward | TCAGTTTGCGGGTGTGCGCAG |
| | Reverse | TTCGTCTCGCCTACACCACCG |
| Tcf3 | Forward | CCGCTTGGACGAGGCCATCC |
| | Reverse | AAGCTCGTGGTCAGTGCGCC |
| Tcf4 | Forward | GGCCGCAGCGCCTTCTCTTTA |
| | Reverse | ACCATCATTGACTCCCCCGAGG |
| β-catenin | Forward | CGCTTGGCTGAACCATCAC |
| | Reverse | GTTCCGCGTCATCCTGATAGT |
| 36B4 | Forward | CGCTTTCTGGAGGGTGTCCGC |
| | Reverse | TGCCAGGACGCGCTTGTACC |
